# Phosphoproteomics implicates glutamatergic and dopaminergic signalling in the antidepressant-like properties of the iron chelator deferiprone

**DOI:** 10.1101/2023.09.06.556022

**Authors:** Volkan Uzungil, Sandra Luza, Carlos M Opazo, Isaline Mees, Shanshan Li, Ching-Seng Ang, Nicholas A Williamson, Ashley I Bush, Anthony J Hannan, Thibault Renoir

## Abstract

Current antidepressants have limitations due to insufficient efficacy and delay before improvement in symptoms. Polymorphisms of the serotonin transporter (5-HTT) gene have been linked to depression (when combined with stressful life events) and to altered response to selective serotonergic reuptake inhibitors. We have previously revealed the antidepressant-like properties of the iron chelator deferiprone in the 5-HTT knock-out (KO) mouse model of depression. Furthermore, deferiprone was found to alter neural activity in the prefrontal cortex of both wild-type (WT) and 5-HTT KO mice. In the current study, we examined the molecular effects of acute deferiprone treatment in the prefrontal cortex of both genotypes via phosphoproteomics. In WT mice treated with deferiprone, there were 22 differentially expressed phosphosites, with gene ontology analysis implicating cytoskeletal proteins. In 5-HTT KO mice treated with deferiprone, we found 33 differentially expressed phosphosites. Gene ontology analyses revealed phosphoproteins that were predominantly involved in synaptic and glutamatergic signalling. In a drug naive cohort, the analysis revealed 21 differentially expressed phosphosites in 5-HTT KO compared to WT mice. We confirmed the deferiprone-induced increase in Tyrosine hydroxylase serine 40 residue phosphorylation (pTH-Ser40) (initially revealed in our phosphoproteomics study) by western blots, with deferiprone increasing pTH-Ser40 expression in WT and 5-HTT KO mice. As glutamatergic and synaptic signalling are dysfunctional in 5-HTT KO mice (and are the target of fast-acting antidepressant drugs such as ketamine), these molecular effects may underpin deferiprone’s antidepressant-like properties. Furthermore, dopaminergic signalling may also be involved in deferiprone’s antidepressant-like properties.

## 1. Introduction

Currently prescribed antidepressants have limited utility due to suboptimal efficacy and a latency period of up to 8 weeks before substantial benefits are observed (Gaynes et al., 2009; Trivedi et al., 2006). Although the new class of fast-acting glutamatergic antidepressants such as ketamine have shown therapeutic potential, they have extensive side-effect profiles (Driesen et al., 2013; Ho & Zhang, 2016; Shiroma et al., 2020). These constraints indicate the need to develop novel fast acting and effective antidepressant therapies.

We have previously discovered that the iron chelator deferiprone has acute antidepressant-like properties on the serotonin transporter knock-out (5-HTT KO) mouse model of depression (Uzungil et al., 2022). Importantly, iron is a co-factor for monoamine neurotransmitter synthesis (Roberts & Fitzpatrick, 2013) and involved in reactive oxygen species production (Crichton et al., 2011), which is implicated in major depressive disorder (MDD) pathophysiology (Bakunina et al., 2015; Bhatt et al., 2020). Critically, we found that acute deferiprone treatment had no effect on levels of iron in multiple brain regions, indicating that deferiprone’s antidepressant-like properties are not mediated directly through altering iron levels (Uzungil et al., 2022).

The new class of fast-acting antidepressants including ketamine, exhibit their antidepressant properties by affecting glutamatergic signalling (Zarate et al., 2006). The mechanism of action includes altering AMPA and NMDA receptor activity, in addition to astrocytic recycling of glutamate via the GLT-1 transporter (Aleksandrova et al., 2017; Y. Chen et al., 2022). The downstream effect of ketamine and typical monoaminergic antidepressants is to enhance synaptic plasticity (Abdallah et al., 2016; R. S. Duman et al., 2016; N. Li et al., 2010). Dysfunctional synaptic plasticity is a prominent pathology of MDD and an important mechanism for the therapeutic action of antidepressant drugs (R. S. Duman et al., 2016).

Additionally, glutamatergic receptor signalling is mediated by nNOS activity, which is an enzyme implicated in MDD by affecting synaptic plasticity (Zhou et al., 2018). Excess levels of nitric oxide has been found in patients with MDD and the SSRI paroxetine reduces levels of nitric oxide, via the inhibition of its precursor enzyme nNOS (Dhir & Kulkarni, 2011; Finkel et al., 1996; Lee et al., 2006; Suzuki et al., 2001). nNOS alters synaptic plasticity by interacting directly with postsynaptic density protein PSD-95, which is bound to NMDA receptors (Christopherson et al., 1999). The nNOS-PSD-95-NMDA synaptic complex has developed as a novel target for antidepressant drugs (Doucet et al., 2012). Relevant to the iron chelator deferiprone, the metal influences levels of nitric oxide; and deferiprone was shown to alter eNOS levels in the periphery (Koskenkorva-Frank et al., 2013; Sriwantana et al., 2018). The nitric oxide system also influences 5-HTT function as activity of nNOS results in phosphorylation of 5-HTT via intracellular kinase activity (Chanrion et al., 2007; Garthwaite, 2007; Zhu et al., 2004).

In addition to glutamate, dopaminergic signalling is dysregulated in MDD patients and drugs acting on dopaminergic signalling can augment antidepressant properties of therapeutics (Belujon & Grace, 2017; Kato & Chang, 2013; Thase et al., 2007). Tyrosine hydroxylase is the precursor enzyme involved in the production of dopamine, with phosphorylation of the serine 40 residue involved in mediating the enzyme’s activity (Daubner et al., 1992). Iron is a co-factor involved in the activity of tyrosine hydroxylase, with the iron chelator deferoxamine increasing levels of pTH-Ser40 in cell culture (Connor et al., 2009; Roberts & Fitzpatrick, 2013).

The 5-HTT KO mouse model was used in the current study as polymorphisms of the 5-HTT gene are implicated in MDD and these mice display depression-related behaviours (Caspi et al., 2003; E. A. Duman & Canli, 2015; Kalueff et al., 2010; Uzungil et al., 2022; Wilson et al., 2020). An additional utility of the 5-HTT KO mouse is that it is considered a treatment resistant model, as both allelic polymorphisms of the 5-HTT gene and 5-HTT KO mice have reduced response to selective serotonin reuptake inhibitor (SSRI) drugs (Holmes et al., 2002; Serretti et al., 2007; Wilson et al., 2020). 5-HTT KO mice also have increased expression of oxidative stress, a mechanism known to be perturbed by iron levels (Crichton et al., 2011; Mössner et al., 2002).

Brain-wide regions are implicated in MDD, with prefrontal cortex dysfunction being consistently paired in both preclinical and clinical studies (Pandya et al., 2012; Pizzagalli & Roberts, 2022). Similarly, the 5-HTT KO mouse model exhibits prefrontal cortex dysfunction that is associated with protein and signalling pathway dysregulation (Goodfellow et al., 2014). We had previously observed that acute deferiprone treatment had an effect on the activity of numerous brain regions, including the prefrontal cortex (Uzungil et al., 2022). Due to the relative functional homogeneity of the prefrontal cortex, in the current study we focused on determining the action of deferiprone in this brain region.

One method to determine global molecular actions of deferiprone is to conduct a phosphoproteomics analysis following acute administration of the drug. Phosphorylation or dephosphorylation is a protein post-translational modification event which can be used as an indirect maker of protein activity or inactivity (Ramazi & Zahiri, 2021). Our lab has previously demonstrated a reversal of phosphoproteomic dysregulation in Huntington’s disease mice following environmental enrichment (Mees, Li, et al., 2022).There is evidence that patients with MDD have an altered phosphoproteome, with synaptic proteins being implicated in the differential expression profile (Martins-de-Souza et al., 2012). A phosphoproteomics analysis has also been utilised to determine that synaptic protein phosphorylation is involved in the antidepressant-like properties of ketamine following acute treatment (Xiao et al., 2020). The 5-HTT KO mouse also has altered prefrontal cortex phosphorylation of key proteins (Goodfellow et al., 2014).

Utilising a phosphoproteomics analysis, whose experimental workflow is depicted in Fig. 1, the current study explored the molecular effects of acute deferiprone treatment on the 5-HTT KO mouse model of depression. To determine the drug’s mechanism of action and potentially link it to the behavioural antidepressant-like properties, we assessed protein phosphorylation changes following acute treatment of deferiprone in the prefrontal cortex.

**Fig. 1.**
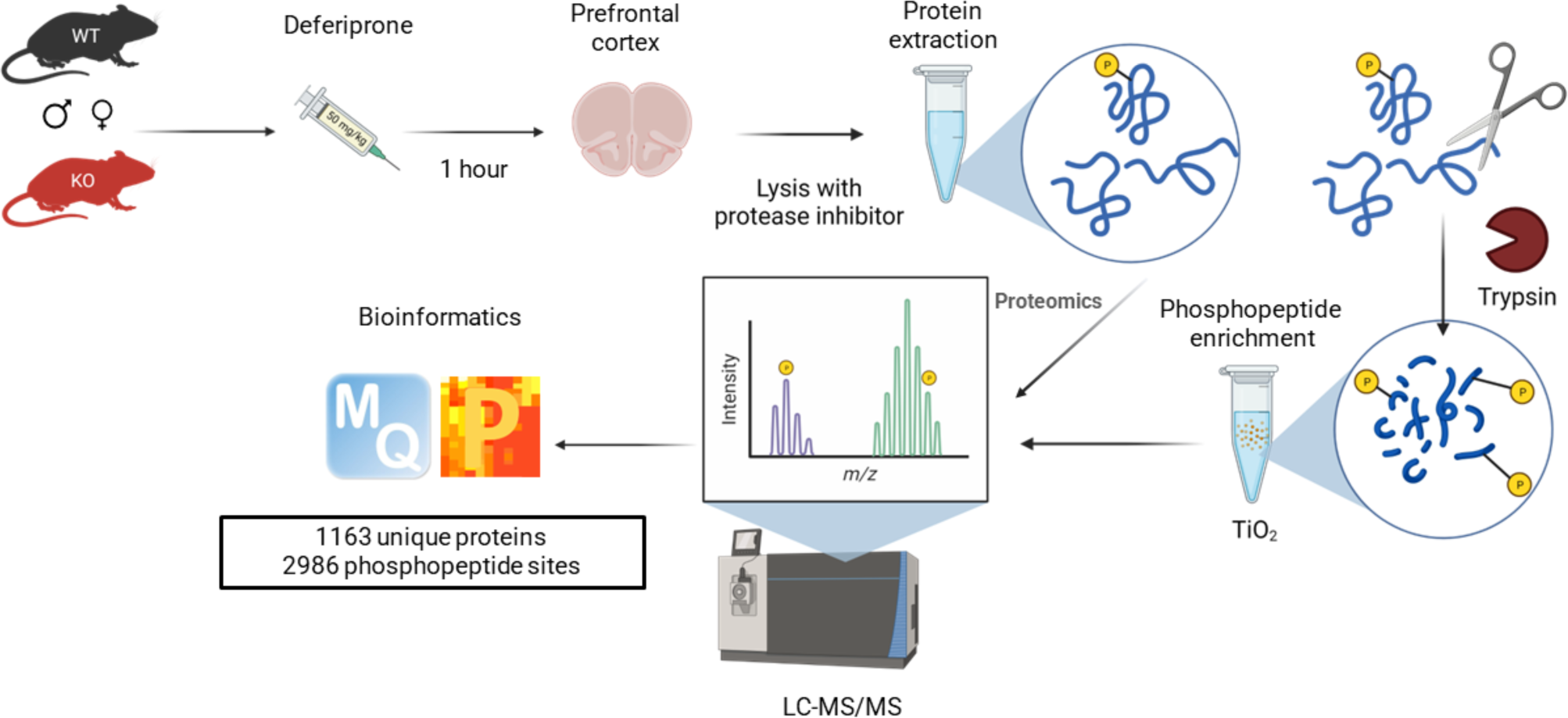
Overview of global proteomics and phosphoproteomics workflow in the prefrontal cortex following acute deferiprone treatment. Created on Biorender.com.

## 2. Materials and methods

### 2.1 Animals and housing

Male and female WT control and 5-HTT KO mice were bred on a C57Bl/6J genetic background from a colony established at the FINMH. Post-weaning, mice were housed (3-5 mice per cage) by littermates in individually ventilated cages until 8 weeks of age. At 8 weeks they were transferred to open top cages (34 x 16 x 16 cm) with standard wood shaving bedding and maintaining original cage-mates. The animals had access to food and water ad libitum while housed in 12/12-hour light/dark cycle rooms (light at 07:00-19:00hrs). All handling or behavioural experiments were done in the light phase. All experiments were conducted between 10-20 weeks of age. All experiments were conducted with the experimenter blind to genotype and treatment. Mice were randomly allocated to treatment groups with even distribution of cage-mates to all groups. All animal care and procedures were approved by the FINMH Animal Ethics Committee and conducted in concordance with guidelines published by the NHMRC.

### 2.2 Pharmacological treatments

The iron chelator 3-Hydroxy-1,2-dimethyl-4(1H)-pyridone (deferiprone; Sigma-Aldrich, Castle Hill, Australia) and 0.9% sterile saline/vehicle (Baxter, Old Toongabbie, Australia) were injected via i.p. administration. Deferiprone was dissolved in 0.9% saline. A clinically relevant dose of 50 mg/kg deferiprone were selected (Hider & Hoffbrand, 2018).

### 2.3 Phosphoproteomics

#### 2.3.1 Sample preparation and phosphopeptide enrichment

Mice were injected with either vehicle or deferiprone and fresh brain tissue was collected 1 hour later, following cervical dislocation. Prefrontal cortex samples from the same hemisphere were used for protein extraction with RIPA buffer (Tris 25 mM, NaCl 150 mM, NP40 1%, sodium deoxycholate 1%, SDS 0.5%) containing protease and phosphatase inhibitors (PhosSTOP, Roche). The lysates were then centrifuged (4°C, 20 minutes, 13000rpm) to remove cell debris. The supernatant was subjected to micro BCA assay (Pierce) where 500ug of proteins were precipitated overnight at - 20°C using 5x volume ice cold acetone. The resultant protein pellet was then solubilized in 8M Urea in 50mM TEAB and incubated for 30 minutes at 37°C. The samples were then treated with TCEP (10mM) for 45 min at 37°C followed by Iodoacetamide (IAM) (55mM) for 30 minutes at room temperature. Thereafter, the samples were diluted to 1M urea with 25mM TEAB and digested overnight at 37°C with Trypsin/LysC (1:50 protein:enzyme). The digested samples were acidified to 1% (v/v) formic acid before solid-phase extraction clean up using Oasis HLB 60mg cartridges. The cartridges were first conditioned with 80% ACN containing 0.1% TFA, followed 0.1% TFA. The samples were then loaded onto the cartridges and washed twice with 0.1% TFA. The proteins were eluted with 80% ACN containing 0.1% TFA and freeze-dried. 20 µg of digested protein was kept for a global unbias proteomics experiment and the remaining peptides were used for phosphopeptide enrichment. For phosphopeptide enrichment, TiO beads (6:1 TiO:peptides) were first washed with 50% ACN, 5% TFA (Washing Buffer) then incubated for 10 minutes with 2M Lactic acid in 5% TFA, 50% ACN (Loading Buffer). The TiO beads in loading buffer were added to the peptides samples and incubated for 1 hour. The phosphopeptides were then washed (twice with the loading buffer and then 3 times with the washing buffer) and sequentially eluted with 1% ammonia and 30% ACN. The samples were then acidified with 1ul of formic acid per 10ul eluent and freeze-dried. The phosphopeptides were resuspended in a buffer containing 2% ACN, 0.05% TFA just before LC-MS/MS analysis.

#### 2.3.2 LC-MS/MS

Samples were analyzed by nanoESI-LC-MS/MS using a Orbitrap Exploris 480 mass spectrometer (Thermo Scientific) equipped with a nanoflow reversed-phase-HPLC (Ultimate 3000 RSLC, Dionex). The LC system was equipped with an Acclaim Pepmap nano-trap column (Dinoex-C18, 100 Å, 75 µm x 2 cm) and an Acclaim Pepmap RSLC analytical column (Dinoex-C18, 100 Å, 75 µm x 50 cm). The tryptic peptides were injected to the enrichment column at an isocratic flow of 5 µL/min of 2% v/v CH3CN containing 0.1% v/v formic acid for 5 min applied before the enrichment column was switched in-line with the analytical column. The eluents were 5% DMSO in 0.1% v/v formic acid (solvent A) and 5% DMSO in 100% v/v CH3CN and 0.1% v/v formic acid (solvent B). The flow gradient was (i) 0-6min at 3% B, (ii) 6-95 min, 3-22% B (iii) 95-105min 22-40% B (iv) 105-110min, 40-80% B (v) 110-115min, 80-80% B (vi) 115-117min, 80-3% and equilibrated at 3% B for 10 minutes before the next sample injection. All spectra were acquired in positive ionization mode with full scan MS acquired from m/z 300-1600 in the FT mode at a mass resolving power of 120 000, after accumulating to an AGC target value of 3.0e6, with a maximum accumulation time of 25 ms. The RunStart EASY-IC lock internal lockmass was used. Data-dependent HCD MS/MS of charge states > 1 was performed using a 3 sec scan method, at a normalized AGC target of 100%, automatic injection, a normalized collision energy of 30%, and spectra acquired at a resolving power of 15000. Dynamic exclusion was used for 20 s.

Raw files were processed in MaxQuant with the Andromeda search engine using default settings for protein and peptide identification. Cysteine carbamidomethyl was selected as fixed modification and methionine oxidation, serine, threonine, and tyrosine phosphorylation as variable modifications. The match between run option was selected. Protein and peptides were filtered to a false discover rate (FDR) of <0.01.

### 2.4 Bioinformatics and statistical analysis

#### 2.4.1 Phosphoproteomics

The processed data was analysed with Perseus (version 1.6.14.0). First, we removed contaminants and reverse peptides from the matrix. Peptides with a phosphate localization probability higher than 0.75 were kept for further analysis. After expanding the site table, phosphopeptides intensities were log2 transformed. Only phosphopeptides with valid values in 100% of the samples in at least one group were kept for statistical analysis. Following this, we normalized the intensities of the phosphopeptides in each sample by subtracting the median and imputed missing values from normal distribution (0.3 width, 1.8 down shift). 2 samples were removed due to being outliers as they were separated from samples in the same group on PCA plot. Partial least squares were used to correct for batch effects using the R package PLSDA-batch (Wang & Lê Cao, 2023). We then performed T-tests comparing 2 groups at a time: p-value 0.01 and fold-change (FC) 1.5 was used as significance. No adjustments were made for multiple comparisons to ensure that true positives were recognised by the analyses and to limit type II errors (Martins-de-Souza et al., 2012; Rothman, 1990). No sex differences were observed and therefore all data was pooled for sex.

The differential expressed phosphopeptides were matched to their gene and from there a gene ontology (GO) enrichment analysis was conducted. The protein class of the differentially expressed phosphoproteins was run on the Panther database (Mi et al., 2021). The phosphoprotein list was then compared against a reference genome to determine the overrepresentation of GO terms using Fisher’s exact test on Panther (Mi et al., 2021). The phosphosite residue associated with each differentially expressed phosphopeptide was then run on PhosphoSitePlus to determine if upstream/downstream regulation of the residue had been previously annotated (Hornbeck et al., 2015). To determine pathways involved in the differentially expressed list, the genes associated with the differential expressed phosphoproteins were analysed using the Kyoto Encyclopedia of Genes and Genomes (KEGG) (Kanehisa et al., 2014). Statistical analysis and visualisations were created with Perseus (version 1.6.14.0) and GraphPad prism 9.0 (Graphpad software. Inc, LA Jolla, USA). Size of treatment group numbers were based on prior studies (n=4-6) for a total of 21 samples (Mees, Li, et al., 2022; Mees, Tran, et al., 2022; Xiao et al., 2020).

#### 2.4.2 Global proteomics

The processed data was analysed with Perseus (version 1.6.14.0). First, we removed contaminants, reverse, and only identified by site peptides from the matrix. The data was then log2 transformed and filtered by valid values, with a minimum of 100% values in at least one of the treatment groups. The intensities of peptides in each sample were normalized by subtracting the median and missing values were imputed from normal distribution (0.3 width, 1.8 down shift). 2 samples were removed due to being outliers as determined by phosphoproteomics analysis. No sex differences were observed and therefore all data was pooled for sex. We then performed two sample T-tests and significant changes were defined as p<0.01 and, Fold change of 1.5. Size of treatment group numbers were based on prior studies (n=5-6) for a total of 23 samples (Mees, Li, et al., 2022; Mees, Tran, et al., 2022; Xiao et al., 2020). Code for phosphoproteomics and global proteomics analysis can be found at https://github.com/vuzungil01/dfp_phosphoproteomics.

### 2.5 Western blot

#### 2.5.1 Tissue collection and processing

Mice were injected with either vehicle or deferiprone and 1 hour later prefrontal cortex tissue was collected following cervical dislocation. Mice were randomly allocated to treatment groups with even distribution of cage-mates to all groups. Tissue was homogenized 1/10 (w/v) in 50 mM Tris-HCl pH 7.5 containing 1% NP-40 (v/v). Samples were then sonicated at 4°C (10 cycles x 10 sec x 10 rest, high). This was followed by centrifuging samples at 10,000 *g* for 10 mins at 4°C. Supernatants were separated from pellets and protein concentrations were determined using BCA Protein Assay Kit (Pierce, USA). Supernatants were aliquoted and kept at −80 °C for protein quantification.

#### 2.5.2 Protein quantification

To detect protein quantity NP-40-soluble fractions (40 µg) of each sample were resolved by electrophoresis in 4-12% SDS-polyacrylamide gels for 1 h at 120 V. Proteins were transferred to a PVDF membrane for 1 h at 15 V. Blots were blocked with 5% (w/v) skim milk in Tris-buffered saline containing 0.1% Tween-20 (TBST). Blots were then incubated overnight with the primary antibodies against pTH-Ser40 (anti-rabbit; Cell Signalling Technology; CAT#2791) diluted in TBST containing 3% (w/v) BSA at 4°C. After four washings with TBST, the blots were incubated with conjugated secondary HRP antibody in TBST, 0.1% SDS for 1h at 22⁰C. Then, the blots were washed four times with TBST for 5 mins each. The blots were finally immersed in Chemiluminescence (ECL) reagents, and developed signals for pTH-Ser40 were quantified using the multi-gauge (V3.0) software. The blots were stripped in glycine stripping buffer (pH=2.2), re-blocking, and tested with a primary antibody against TH (anti-rabbit; Merck Millipore; CAT#AB152), repeating the same procedures to obtain and quantify the TH signals. Internal controls were made with 5 mL from all samples and was used to normalize the signals between gels.

#### 2.5.3 Statistical analysis

The levels of pTH-Ser40 and TH were calculated as the average of duplicate gels with the exception of samples used in pilot studies. No sex differences were found and therefore all data was pooled for sex. The analyses were performed with a non-parametric Kruskal-Wallis test with a Dunnett’s post-hoc test for comparing individual groups Significance was set at p<0.05 and a statistically valid trend was set at 0.05<p<0.07. Treatment group sizes (n=10-14) were extrapolated from prior western blot studies to conduct sex differences statistical analysis (Wright et al., 2016). This resulted in a total of 47 samples used for this study. Statistical analysis and visualisation were conducted using GraphPad prism 9.0 (Graphpad software. Inc, LA Jolla, USA).

## 3. Results

To determine the molecular actions of acute deferiprone treatment, the phosphoproteome of WT and 5-HTT KO mice were determined in the prefrontal cortex following delivery of the drug. Following phosphoproteomics enrichment, 2986 phosphopeptide sites and 1163 unique proteins were identified. To ensure that variability between biological samples within each group was low we determined Pearson’s correlation (r) for WT-VEH=0.9296, WT-DFP=0.9176, KO-VEH=0.9338 and KO-DFP=0.9730.

### 3.1 Cytoskeletal signalling protein phosphorylation differentially expressed in WT mice following acute deferiprone treatment

The phosphoproteomics analysis revealed 2 upregulated and 20 downregulated phosphosites in WT mice treated with deferiprone (Fig. 2A). A table of the upregulated and downregulated phosphosites with their associated biological relevance was compiled (Table 1).

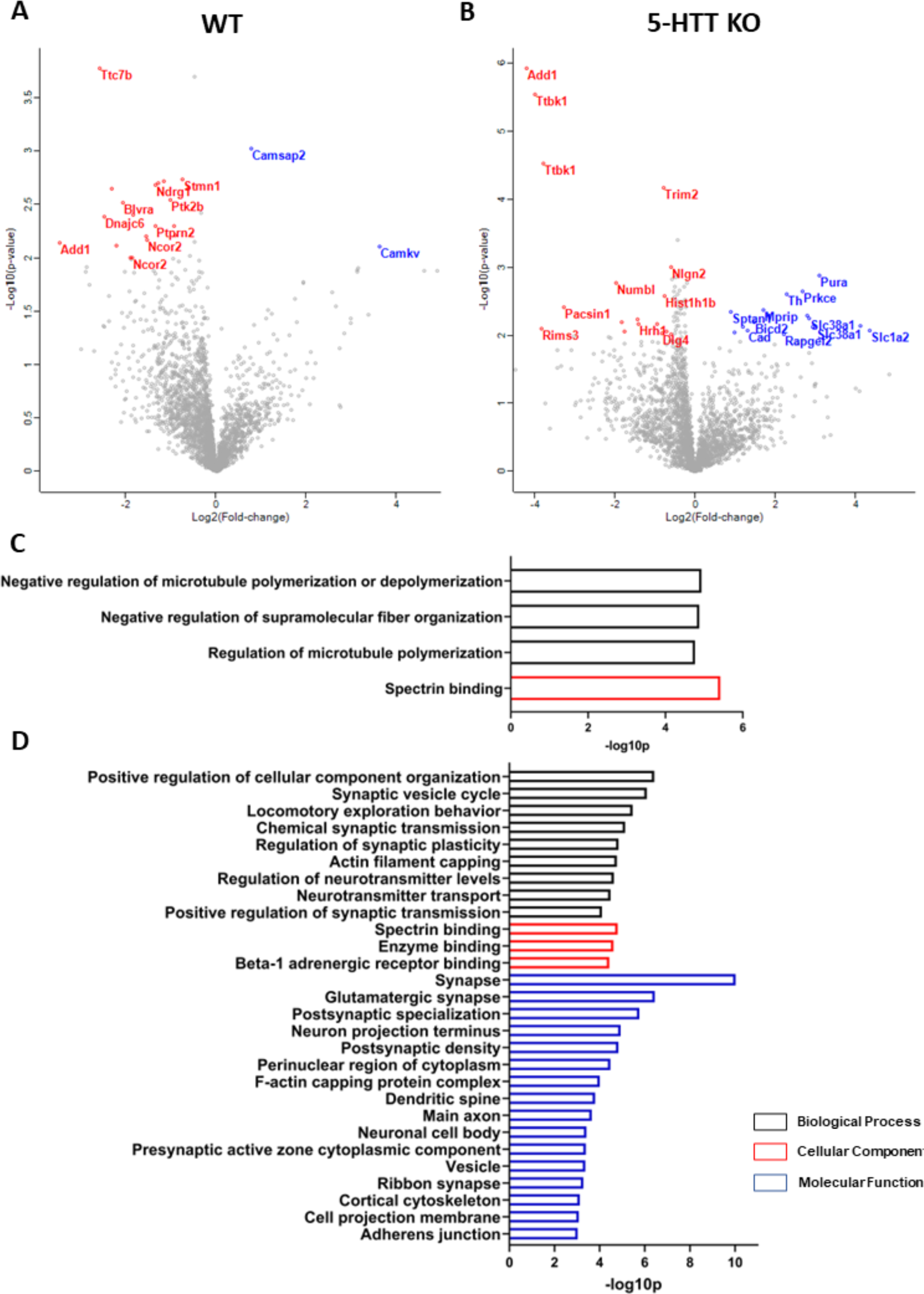
Phosphoproteome of WT and 5-HTT KO mice following deferiprone treatment in the prefrontal cortex. (A) Volcano plot revealed 2 upregulated phosphosites in WT mice treated with deferiprone and with 20 downregulated phosphosites in comparison to vehicle controls. Each dot represents a unique phosphosite with the differentially expressed highlighted; blue (up-regulated) and red (down-regulated). (B) Volcano plot revealed 17 upregulated phosphosites in KO mice treated with deferiprone and with 16 downregulated phosphosites in comparison to vehicle controls. Each dot represents a unique phosphosite with the differentially expressed highlighted. A gene ontology overrepresentation analysis against a reference genome of the differentially expressed phosphosites revealed a different profile in WT (C) and KO (D) mice treated with deferiprone. Black bars represent biological process, red bars represent molecular function and blue bars represent cellular component. n=4-6 mice per group.

**Table 1.**
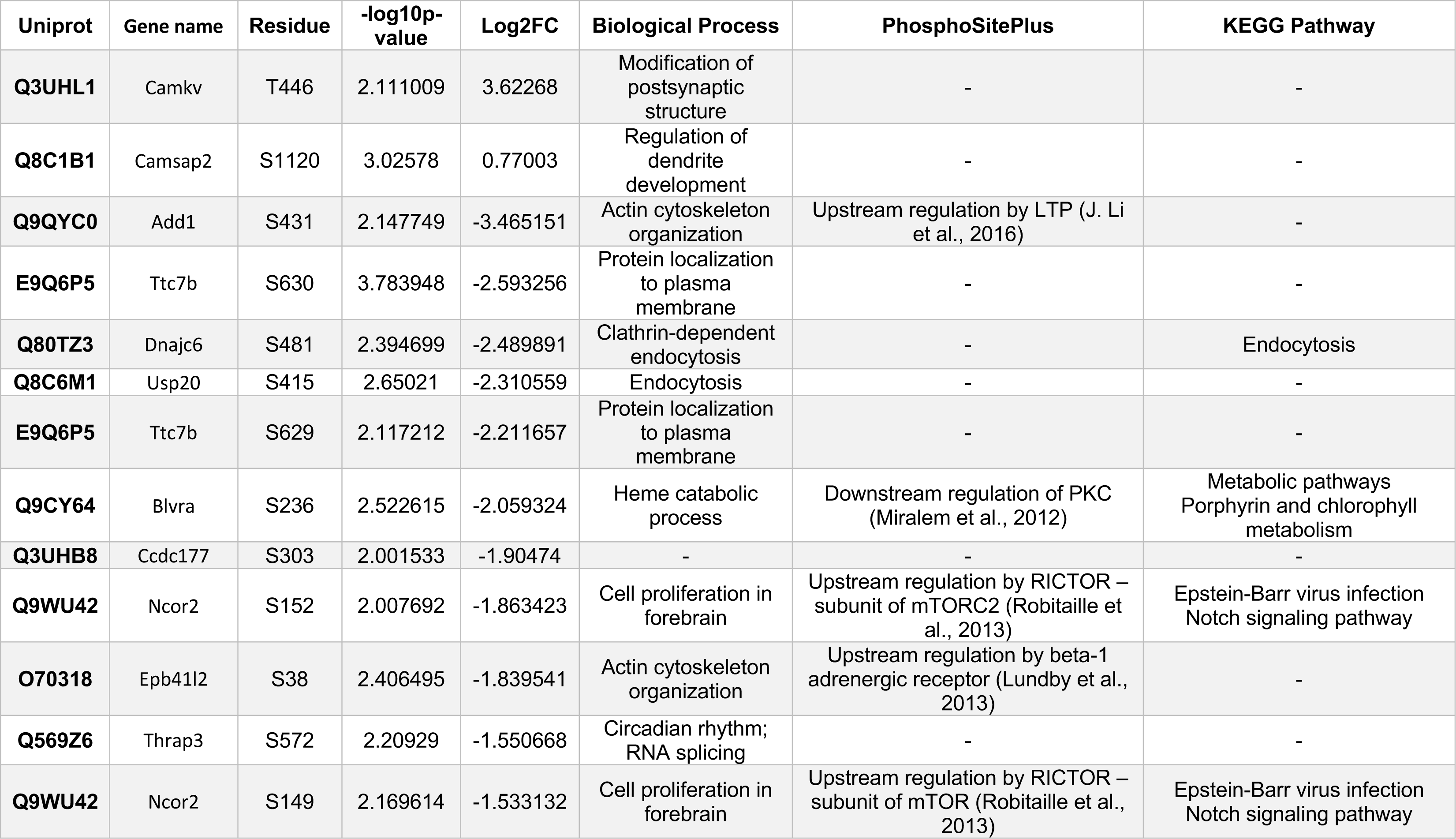
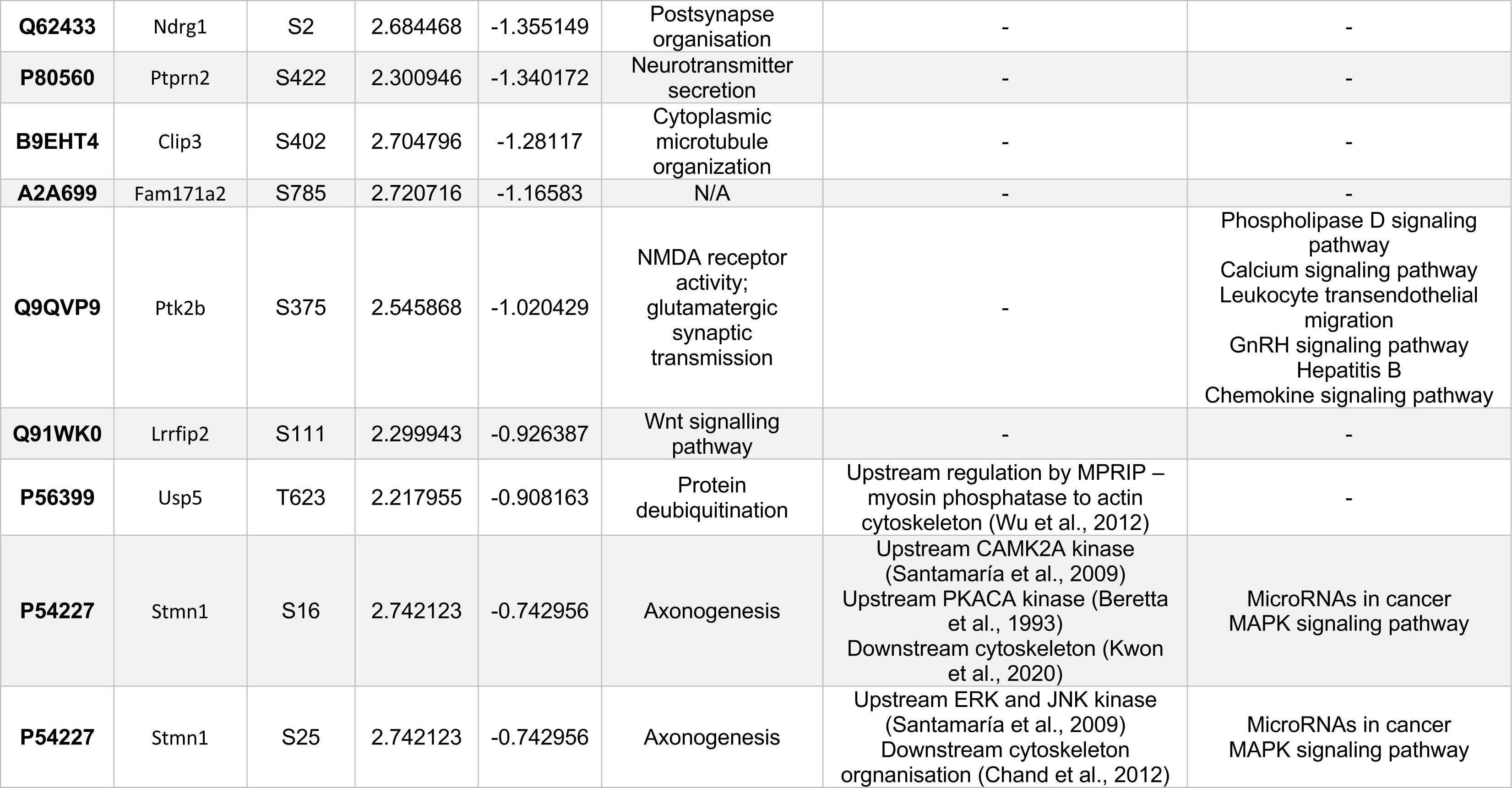
Differentially expressed phosphosites in WT mice treated with deferiprone and relevant biological process, upstream/downstream regulation and KEGG pathway involvement. Fold changes are represented in the Log2 scale (Log2FC).

Of the 22 differentially expressed phosphosites a gene ontology overrepresentation analysis revealed that the equivalent genes involved in cytoskeletal organisation were implicated (Fig. 2C). These included the biological process “regulation of microtubule organisation” (Stmn1, Camsap2, Clip3) and molecular function “spectrin binding” (Add1, Camsap2, Epb41l2).

To determine the relevance of the phosphorylation of the protein residue in upstream or downstream pathways each differentially expressed phosphosite was compared against the PhosphoSitePlus database (Table 1). Of the downregulated phosphosites, upstream kinases which were shown to previously phosphorylate certain protein residues included mTOR (pNcor2-Ser152 & pNcor2-Ser149), CAMK2A (pStmn1-Ser16), PKACA (pStmn1-Ser16), ERK (pStmn1-Ser25) & JNK (pStmn1-Ser25). Additionally, LTP (pAdd1-Ser431) and beta-1 adrenergic receptor activity (pEpb41l2-Ser38) were shown to be upstream regulators. In terms of downstream regulation, there was one identified regulator of PKC activity (pBlvra-Ser236). The analysis also revealed prior annotation of cytoskeletal organisation, as pUsp5-Tyr623 was shown to be upstream regulated by MPRIP while pStmn1-Ser16 and pStmn1-Ser25 were both shown to be downstream regulators of cytoskeletal organisation.

### 3.2 Synaptic, cytoskeletal and glutamatergic signalling protein phosphorylation differentially expressed in 5-HTT KO mice following acute deferiprone treatment

The phosphoproteomics analysis revealed 17 upregulated and 16 downregulated phosphosites in 5-HTT KO mice treated with deferiprone (Fig. 2B). A table of the upregulated (Table 2) and downregulated (Table 3) phosphosites with their associated biological relevance was compiled.

**Table 2.**
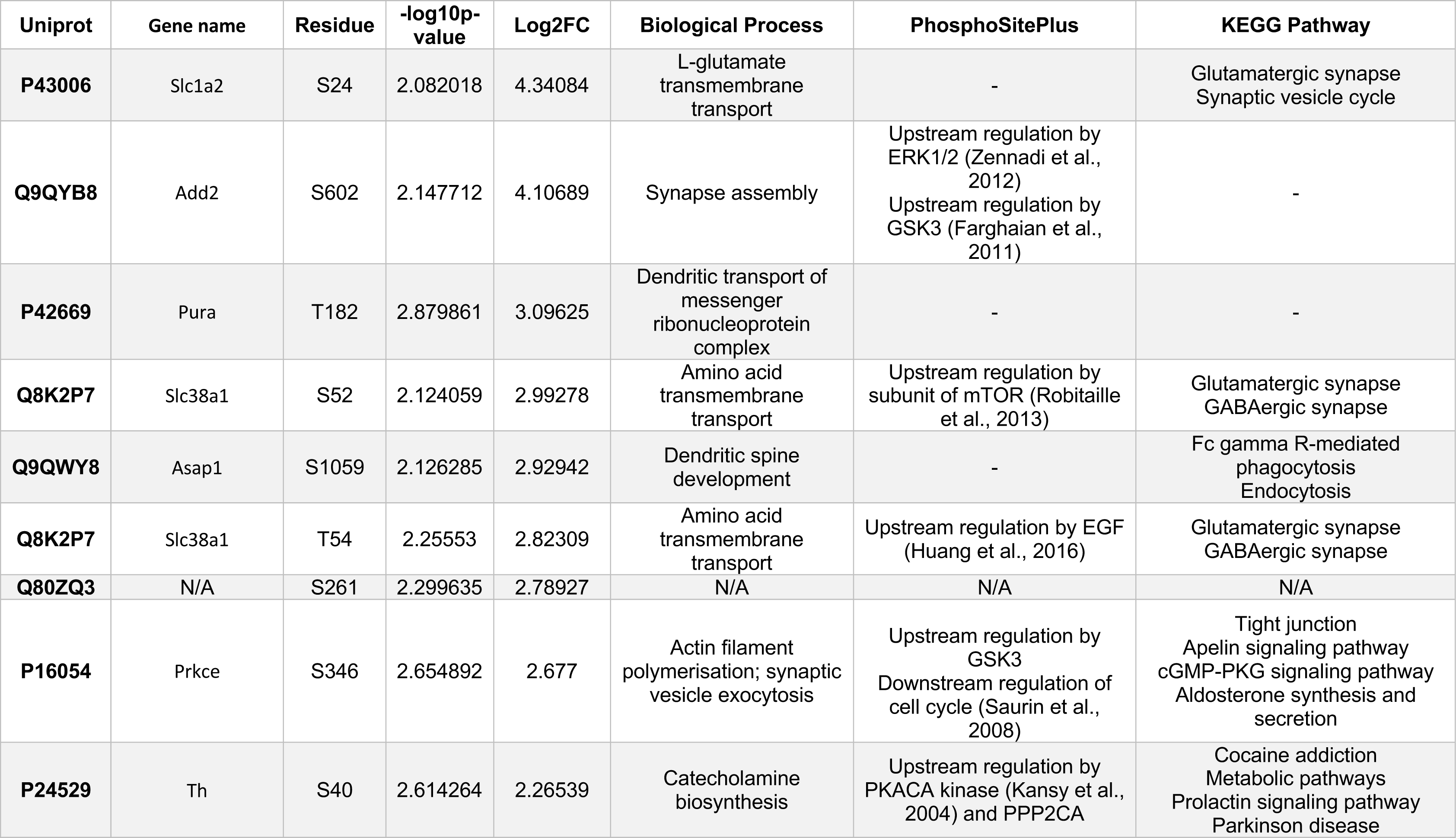
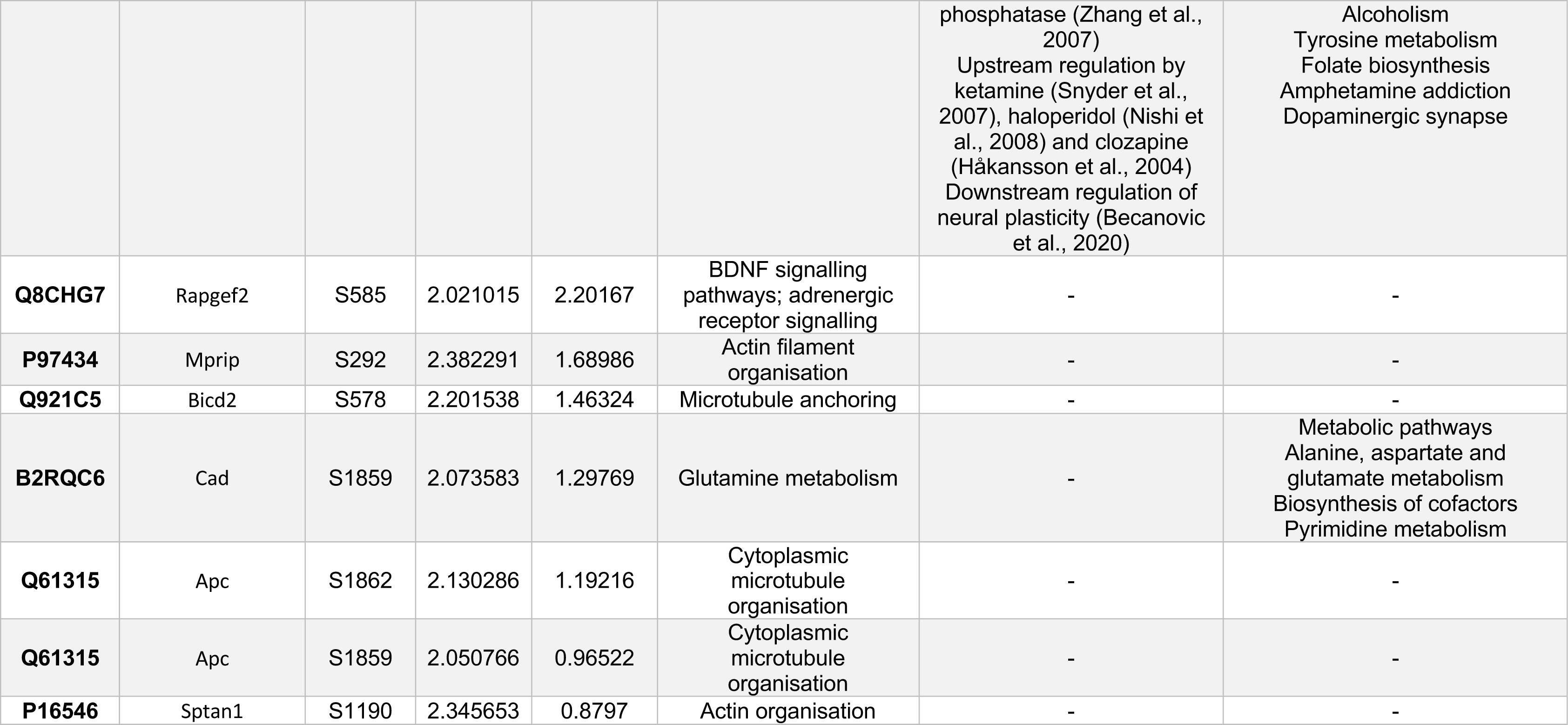
Upregulated phosphosites in 5-HTT KO mice treated with deferiprone and relevant biological process, upstream/downstream regulation and KEGG pathway involvement. Fold changes are represented in the Log2 scale (Log2FC).

**Table 3.**
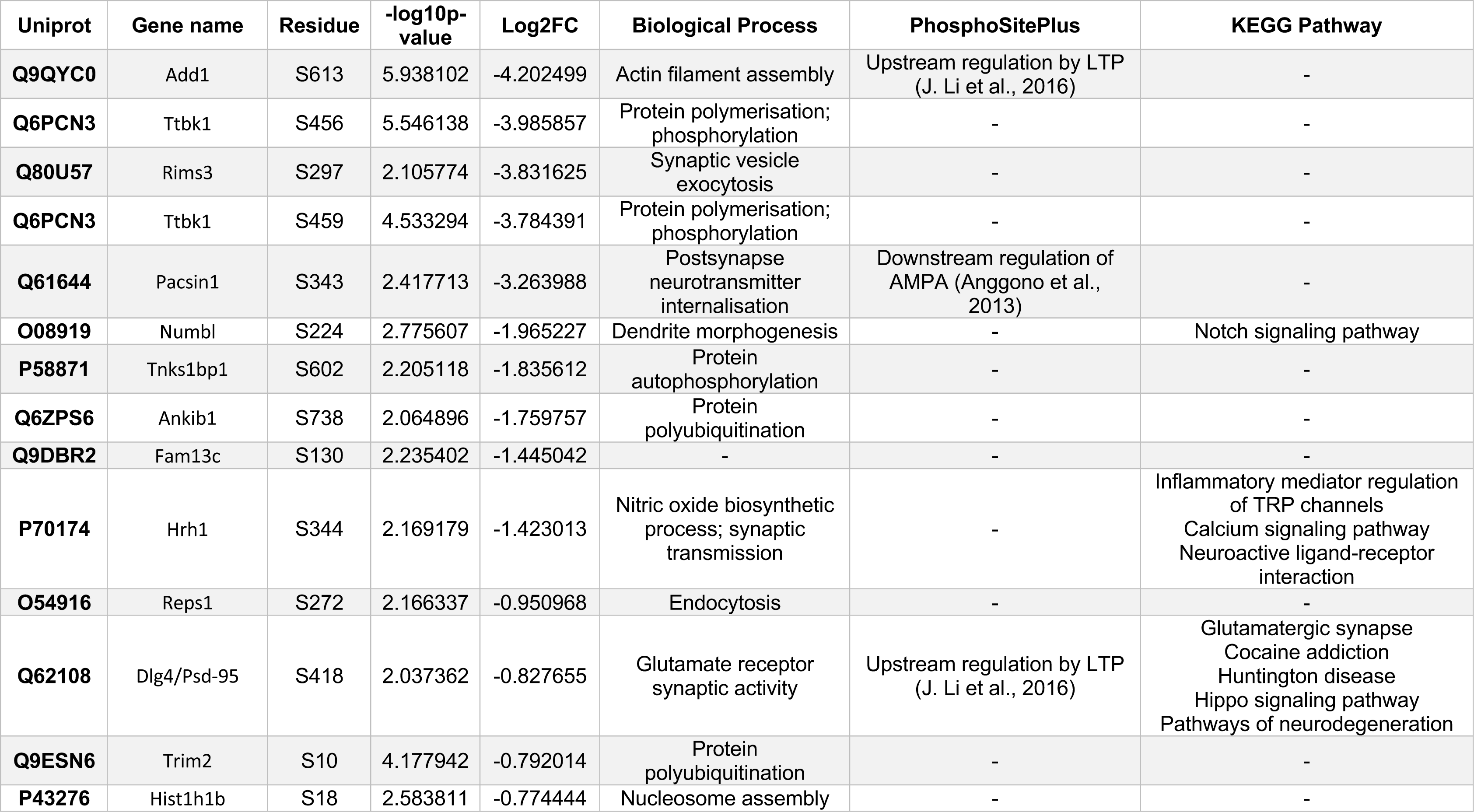
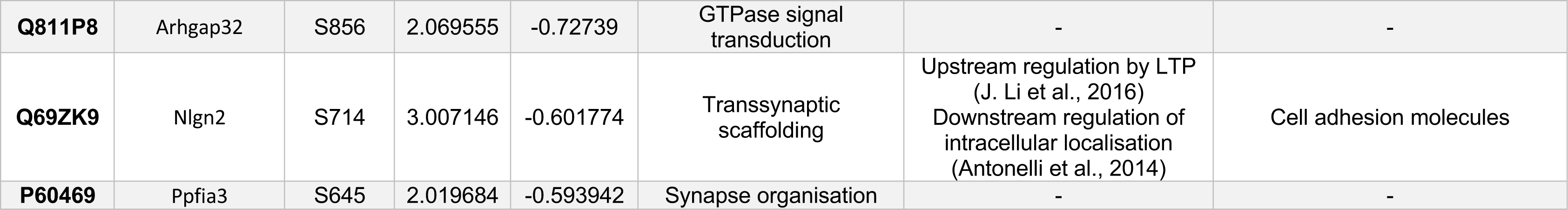
Downregulated phosphosites in 5-HTT KO mice treated with deferiprone and relevant biological process, upstream/downstream regulation and KEGG pathway involvement. Fold changes are represented in the Log2 scale (Log2FC).

| Uniprot | Gene name | Residue | -log10p-value | Log2FC | Biological Process | PhosphoSitePlus | KEGG Pathway |
| --- | --- | --- | --- | --- | --- | --- | --- |
| **Q9QYC0** | Add1 | S613 | 5.938102 | -4.202499 | Actin filament assembly | Upstream regulation by LTP (J. Li et al., 2016) | - |
| **Q6PCN3** | Ttbk1 | S456 | 5.546138 | -3.985857 | Protein polymerisation; phosphorylation | - | - |
| **Q80U57** | Rims3 | S297 | 2.105774 | -3.831625 | Synaptic vesicle exocytosis | - | - |
| **Q6PCN3** | Ttbk1 | S459 | 4.533294 | -3.784391 | Protein polymerisation; phosphorylation | - | - |
| **Q61644** | Pacsin1 | S343 | 2.417713 | -3.263988 | Postsynapse neurotransmitter internalisation | Downstream regulation of AMPA (Anggono et al., 2013) | - |
| **O08919** | Numbl | S224 | 2.775607 | -1.965227 | Dendrite morphogenesis | - | Notch signaling pathway |
| **P58871** | Tnks1bp1 | S602 | 2.205118 | -1.835612 | Protein autophosphorylation | - | - |
| **Q6ZPS6** | Ankib1 | S738 | 2.064896 | -1.759757 | Protein polyubiquitination | - | - |
| **Q9DBR2** | Fam13c | S130 | 2.235402 | -1.445042 | - | - | - |
| **P70174** | Hrh1 | S344 | 2.169179 | -1.423013 | Nitric oxide biosynthetic process; synaptic transmission | - | Inflammatory mediator regulation of TRP channels<br>Calcium signaling pathway<br>Neuroactive ligand-receptor interaction |
| **O54916** | Reps1 | S272 | 2.166337 | -0.950968 | Endocytosis | - | - |
| **Q62108** | Dlg4/Psd-95 | S418 | 2.037362 | -0.827655 | Glutamate receptor synaptic activity | Upstream regulation by LTP (J. Li et al., 2016) | Glutamatergic synapse<br>Cocaine addiction<br>Huntington disease<br>Hippo signaling pathway<br>Pathways of neurodegeneration |
| **Q9ESN6** | Trim2 | S10 | 4.177942 | -0.792014 | Protein polyubiquitination | - | - |
| **P43276** | Hist1h1b | S18 | 2.583811 | -0.774444 | Nucleosome assembly | - | - |
| **Q811P8** | Arhgap32 | S856 | 2.069555 | -0.72739 | GTPase signal transduction | - | - |
| **Q69ZK9** | Nlgn2 | S714 | 3.007146 | -0.601774 | Transsynaptic scaffolding | Upstream regulation by LTP (J. Li et al., 2016)<br>Downstream regulation of intracellular localisation (Antonelli et al., 2014) | Cell adhesion molecules |
| **P60469** | Ppfia3 | S645 | 2.019684 | -0.593942 | Synapse organisation | - | - |

Of the 33 differentially expressed phosphosites, a gene ontology overrepresentation analysis revealed that the equivalent genes involved in synaptic signalling and cytoskeletal organisation were implicated (Fig. 2D). These included biological processes such as “synaptic vesicle cycle” (Ppfia3, Rims3, Nlgn2, Th, Pacsin1) and “positive regulation of cellular component organisation” (Dlg4, Add1, Numbl, Nlgn2, Ttbk1, Rapgef2, Prkce, Asap1, Pacsin1, Apc). Cyotskeletal organisation was also reflected in the molecular function such as “spectrin binding” (Add1, Add2, Sptan1) while synaptic proteins were reflected in the cellular component analysis such as “synapse” (Ppfia3, Dlg4, Add1, Add2, Rims3, Nlgn2, Pura, Th, Rapgef2, Prkce, Asap1, Pacsin1, Arhgap32, Slc1a2, Apc).

To determine the relevance of the phosphorylation of the protein residue in upstream or downstream pathways each differentially expressed phosphosite was compared against the PhosphoSitePlus database (Table 2 & 3). Of the upregulated phosphosites, upstream kinases which were shown to previously phosphorylate these phosphorylation sites included ERK1/2 (pAdd2-Ser602), GSK3 (pAdd2-Ser602 & pPrkce-Ser346), mTOR (pSlc38a1-Ser52) and PKACA (pTh-Ser40). Psychoactive drugs such as ketamine, haloperidol and clozapine were shown to also be upstream regulators for pTH-Ser40 with downstream regulation of neural plasticity. In terms of the downregulated phosphosites, long-term potentiation (LTP) was shown to be upstream regulator of pAdd1-Ser613 and pNlgn2-Ser714, as well as pPacsin1-Ser343 a downstream regulator of the glutamatergic AMPA receptor.

Finally, of the differentially expressed phosphoproteins, a KEGG pathway analysis was conducted to determine which biological pathway the protein was previously implicated in (Table 2 & 3). The only KEGG pathway which had multiple hits was “glutamatergic synapse” (Slc1a2, Slc38a1, Dlg4). Related to glutamatergic synapse was also the KEGG pathway “alanine, aspartate and glutamate metabolism” (Cad). Meanwhile other KEGG pathways implicated in brain processes include “synaptic vesicle cycle” (Slc1a2), “GABAergic synapse” (Slc38a1), “dopaminergic synapse” (Th) and “neuroactive ligand-receptor interaction” (Hrh1).

### 3.3 Gene ontology analysis reveals no significant pathway implicated in the change to phosphoproteome due to genetic ablation of 5-HTT

The phosphoproteomics analysis revealed 11 upregulated and 10 downregulated phosphosites in 5-HTT KO compared to WT mice (Fig. 3A). A table of the upregulated and downregulated phosphosites with their associated biological relevance was compiled (Table 4).

**Fig. 3.**
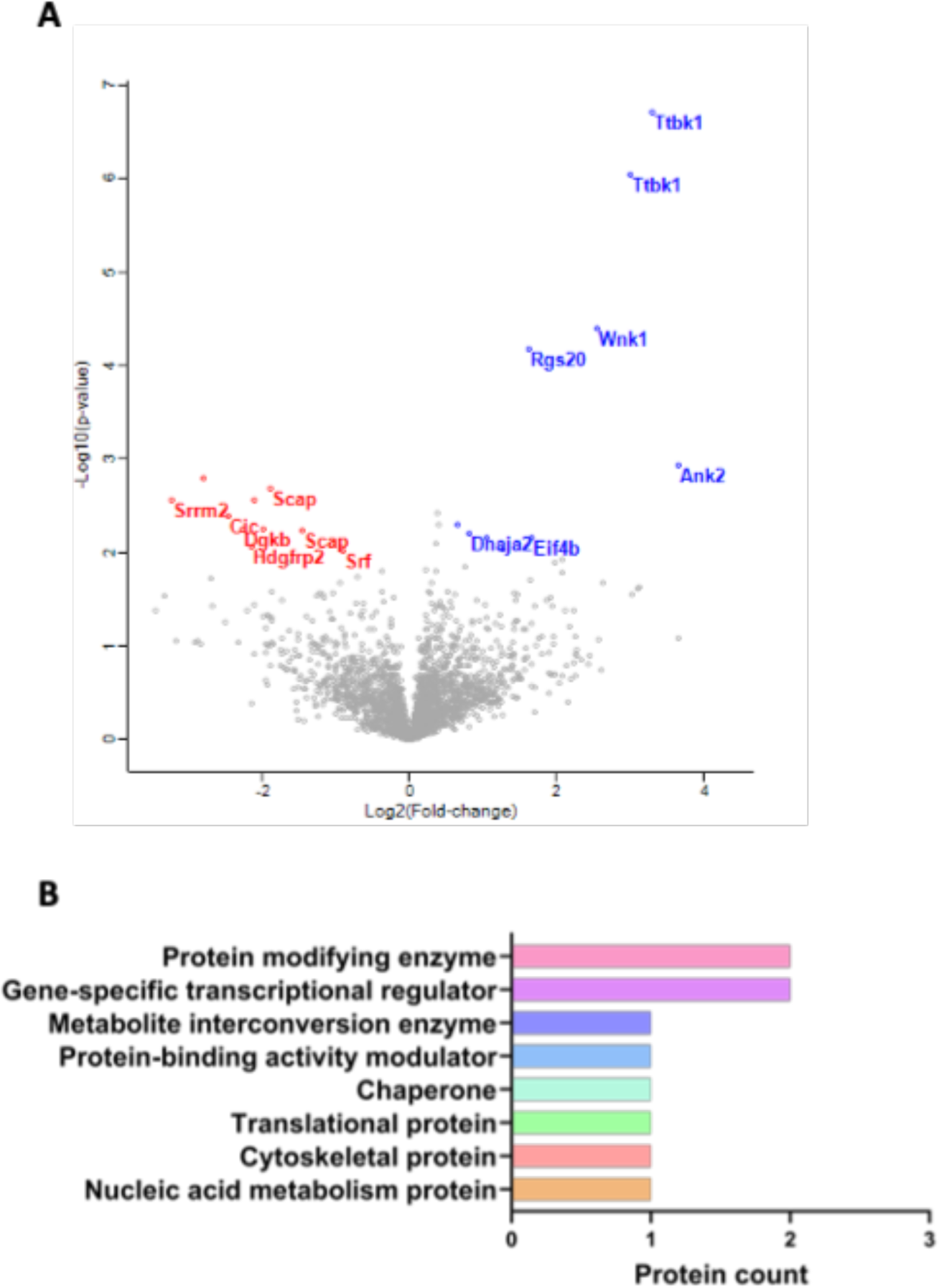
Phosphoproteome profile of treatment naive 5-HTT KO mice compared to WT controls in the prefrontal cortex. (A) Volcano plot revealed 11 upregulated phosphopeptides in the 5-HTT KO mice with 10 downregulated phosphopeptides in comparison to WT controls. Each dot represents a unique phosphopeptide with the differentially expressed highlighted; blue (up-regulated) and red (down-regulated). (B) The gene ontology analysis identified the protein class for differentially expressed phosphopeptides with “protein modifying enzyme” and “gene-specific transcriptional regulator” the only hits with multiple proteins in 5-HTT KO mice compared to WT controls. n=5-6 animals per group.

**Table 4.** Differentially expressed phosphosites in 5-HTT KO mice compared to WT controls in the prefrontal cortex with relevant biological process, upstream/downstream regulation and KEGG pathway involvement. Fold changes are represented in the Log2 scale (Log2FC).

| Uniprot | Gene name | Protein name | Residue | -log10p-value | Log2FC | Biological Process | PhosphoSitePlus | KEGG Pathway |
| --- | --- | --- | --- | --- | --- | --- | --- | --- |
| **Q8C8R3** | Ank2 | Ankyrin-2 | S3781 | 2.940138 | 3.65765 | Calcium ion transporter | Upstream regulation by LTP (J. Li et al. 2016) | Proteoglycans in cancer |
| **Q6PCN3** | Ttbk1 | Tau-tubulin kinase 1 | S456 | 6.723777 | 3.30476 | Protein polymerisation; phosphorylation | - | - |
| **Q6PCN3** | Ttbk1 | Tau-tubulin kinase 1 | S459 | 6.050869 | 3.0072 | Protein polymerisation; phosphorylation | - | - |
| **P83741** | Wnk1 | Serine/threonine-protein kinase WNK1 | S2027 | 4.399559 | 2.55303 | Sodium ion transport | - | - |
| **Q8BGD9** | Eif4b | Eukaryotic translation initiation factor 4B | S409 | 2.165922 | 1.66079 | Translational initiation | - | Proteoglycans in cancer<br>mTOR signalling pathway<br>PI3K-Akt signalling pathway |
| **Q9QZB1** | Rgs20 | Regulator of G-protein signaling 20 | S15 | 4.179089 | 1.62663 | G protein-receptor signaling | - | - |
| **Q9QZB1** | Rgs20 | Regulator of G-protein signaling 20 | S13 | 2.042038 | 1.2725 | G protein-receptor signaling | - | - |
| **Q6DFV3** | Arhgap21 | Rho GTPase-activating protein 21 | S314 | 2.169595 | 1.04929 | GTPase signal transduction | - | - |
| **Q9QYJ0** | Dnaja2 | DnaJ homolog subfamily A member 2 | S394 | 2.206915 | 0.80753 | Cell proliferation | - | Protein processing in endoplasmic reticulum |
| **Q9QYJ0** | Dnaja2 | DnaJ homolog subfamily A member 2 | S395 | 2.206915 | 0.80753 | Cell proliferation | - | Protein processing in endoplasmic reticulum |
| **Q8BL80** | Arhgap22 | Rho GTPase-activating protein 22 | S397 | 2.302286 | 0.66214 | Postsynapse organization; signal transduction | - | - |
| **Q8BTI8** | Srrm2 | Serine/arginine repetitive matrix protein 2 | S2656 | 2.566881 | -<br>3.232283 | mRNA splicing | - | - |
| **Q8BTI8** | Srrm2 | Serine/arginine repetitive matrix protein 2 | S2660 | 2.802669 | -<br>2.805153 | mRNA splicing | - | - |
| **Q924A2** | Cic | Protein capicua homolog | T2310 | 2.402026 | -<br>2.454973 | Learning; social behaviour | - | Spinocerebellar ataxia |
| **Q6NS52** | Dgkb | Diacylglycerol kinase beta | S95 | 2.264498 | -<br>2.274152 | Chemical synaptic transmission | - | Phospholipase D pathway<br>Glycerolipid metabolism<br>Choline metabolism in cancer<br>Phosphatidylinositol signaling system |
| **Q3UMU9** | Hdgfrp2 | Hepatoma-derived growth factor-related protein 2 | S450 | 2.070752 | -<br>2.150609 | Positive regulation of cell growth | Upstream regulation by RIP3 (Wu et al. 2012) | - |
| **Q8K019** | Bclaf1 | Bcl-2-associated transcription factor 1 | S510 | 2.563508 | -<br>2.117053 | Apoptotic process | - | - |
| **Q9QYR6** | Map1a | Microtubule-associated protein 1A | S1772 | 2.254222 | -<br>1.998184 | Regulation of synaptic plasticity | - | - |
| **Q6GQT6** | Scap | Sterol regulatory element-binding protein cleavage-activating protein | S843 | 2.690433 | -<br>1.898801 | Cellular lipid metabolic process | - | - |
| **Q6GQT6** | Scap | Sterol regulatory element-binding protein cleavage-activating protein | S850 | 2.247987 | -<br>1.459237 | Cellular lipid metabolic process | Upstream regulation by Torin 1 mTor inhibitor (Hsu et al. 2011) | - |
| **Q9JM73** | Srf | Serum response factor | S220 | 2.030841 | -<br>0.911628 | Actin cytoskeleton organization | - | Human T-cell leukemia virus 1 infection<br>MAPK signalling pathway<br>cGMP-PKG signalling pathway |

Of the 21 differentially expressed phosphosites, there was no difference in the gene ontology overrepresentation analysis. Instead, a gene ontology analysis with the associated protein count was determined (Fig. 3B). Gene ontology analysis revealed that “protein modifying enzyme” had 2 associated proteins (Wnk1 and Ttbk1) while the other class with multiple proteins was “gene-specific transcriptional regulator” (Cic and Hdgfrp2). All other classes from the gene ontology analysis had 1 associated protein which included “chaperone” (Dnaja2) and “cytoskeletal protein” (Map1a).

To determine the relevance of the phosphorylation of the protein residue in upstream or downstream pathways each differentially expressed phosphosite was compared against the PhosphoSitePlus database (Table 4). Of the upregulated phosphosites, LTP was shown to be an upstream regulator of pAnk2-Ser3781. Meanwhile of the downregulated phosphosites, there was upstream regulation by RIP3 of pHdgfrp2-Ser450 and by an mTOR inhibitor of pScap-Ser850.

### 3.4 Global proteomics reveals minimal differential expression of proteins in the prefrontal cortex of 5-HTT KO mice or deferiprone treatment

A global proteomics analysis was also conducted of WT and 5-HTT KO mice in the prefrontal cortex of mice following delivery of the drug. The global proteomics analysis identified 2615 unique proteins when there were 100% valid values in at least one group with the differentially expressed proteins in treatment groups shown in Table 5. To ensure that variability between biological samples within each group was low we determined Pearson’s correlation (r) for WT-VEH=0.9679, WT-DFP=0.9559, KO-VEH=0.9698 and KO-DFP=0.9703.

**Table 5.** Differentially expressed proteins of group comparisons in the prefrontal cortex and associated biological process during global proteomics analysis. Fold changes are represented in the Log2 scale (Log2FC).

| Uniprot | Condition | Gene | Protein | -Log10(p-value) | Log2(Fold-change) | Biological Process |
| --- | --- | --- | --- | --- | --- | --- |
| **Q8BJ71** | WT-VEH vs KO-VEH | Nup93 | Nuclear pore complex protein Nup93 | 2.34 | 0.88 | mRNA transport |
| **O70250** | WT-VEH vs KO-VEH | Pgam2 | Phosphoglycerate mutase 2 | 2.96 | -2.99 | Glycolysis |
| **Q91XF0** | WT-VEH vs KO-VEH | Pnpo | Pyridoxine-5-phosphate oxidase | 2.63 | -2.13 | Pyridoxine biosynthesis |
| **Q9CRD2** | WT-VEH vs KO-VEH | Emc2 | ER membrane protein complex subunit 2 | 3.42 | -1.23 | Membrane intertase activity |
| **O70456** | WT-VEH vs WT-DFP | Sfn | 14-3-3 protein sigma | 2.11 | 4.54 | Keratinisation |
| **Q8BJ71** | WT-VEH vs WT-DFP | Nup93 | Nuclear pore complex protein Nup93 | 2.24 | 0.73 | Nuclear envelope organisation |

In each comparison differentially expressed proteins are represented in the volcano plot, including the proteins associated to the differentially expressed phosphosite in each condition (Figure 4). Differentially expressed proteins due to genetic ablation of 5-HTT were Nup93, Pgam2, Pnpo and Emc2. Meanwhile differentially expressed proteins in WT mice treated with deferiprone was Sfn and Nup93 (Table 5).

**Fig. 4.**
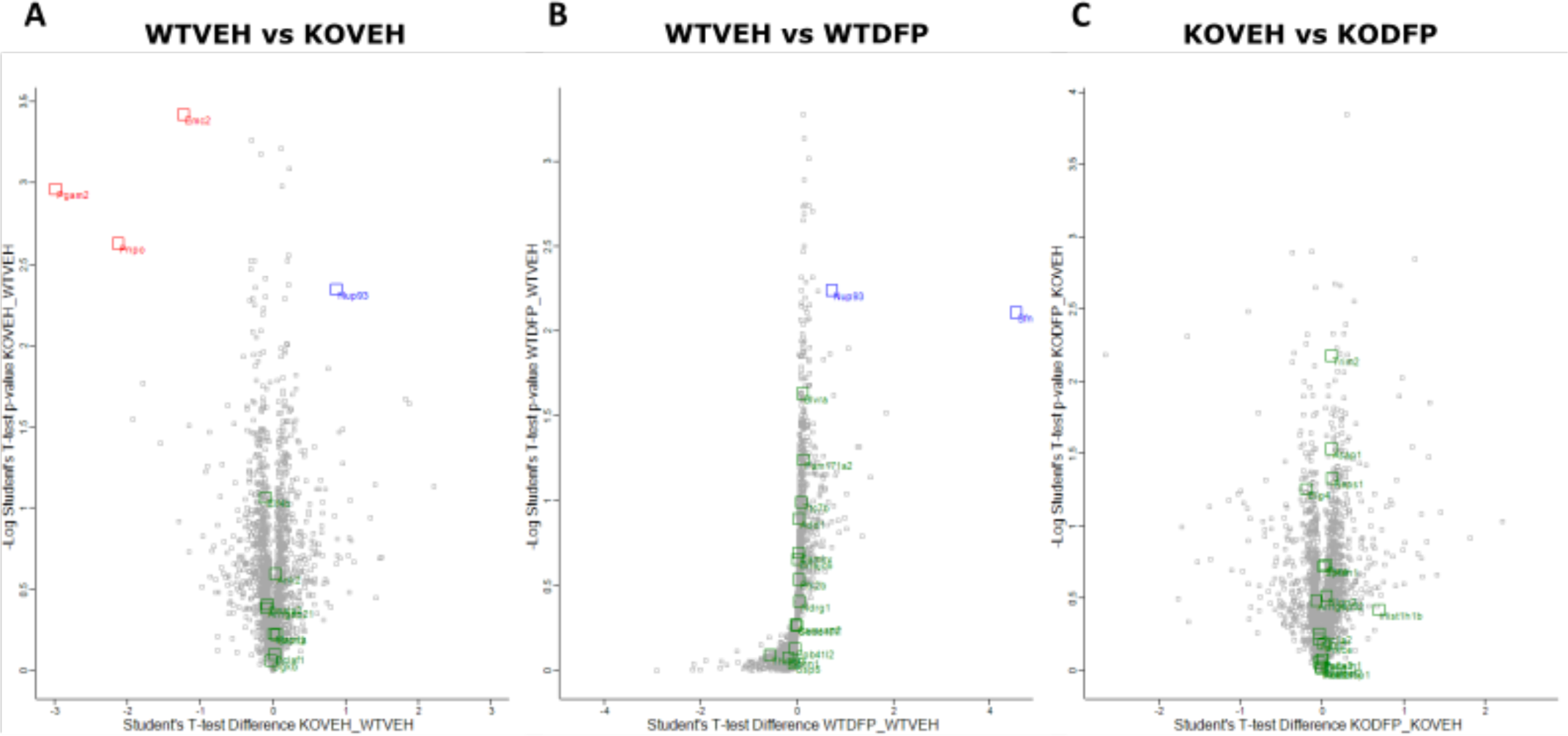
Volcano plot of global proteomics analysis of acute deferiprone treatment in WT and 5-HTT KO mice. There were 4 differentially expressed proteins in 5-HTT KO mice compared to controls (A) and 2 differentially expressed proteins in WT mice treated with deferiprone (B). There were no differentially expressed proteins in 5-HTT KO mice treated with deferiprone (C). Each dot represents a unique phosphosite with the differentially expressed highlighted; blue (up-regulated) and red (down-regulated). Green dots represent proteins associated to differentially expressed phosphosites in phosphoproteomics analysis.

### 3.5 WT and 5-HTT KO mice have increased pTH-Ser40 and ratio of pTH/TH protein levels following acute deferiprone treatment

To confirm findings of increased pTH-Ser40 levels in the phosphoproteomics study, a western blot follow-up analysis was conducted on prefrontal cortex regions following acute deferiprone treatment in WT and 5-HTT KO mice (Figure 5).

**Fig. 5.**
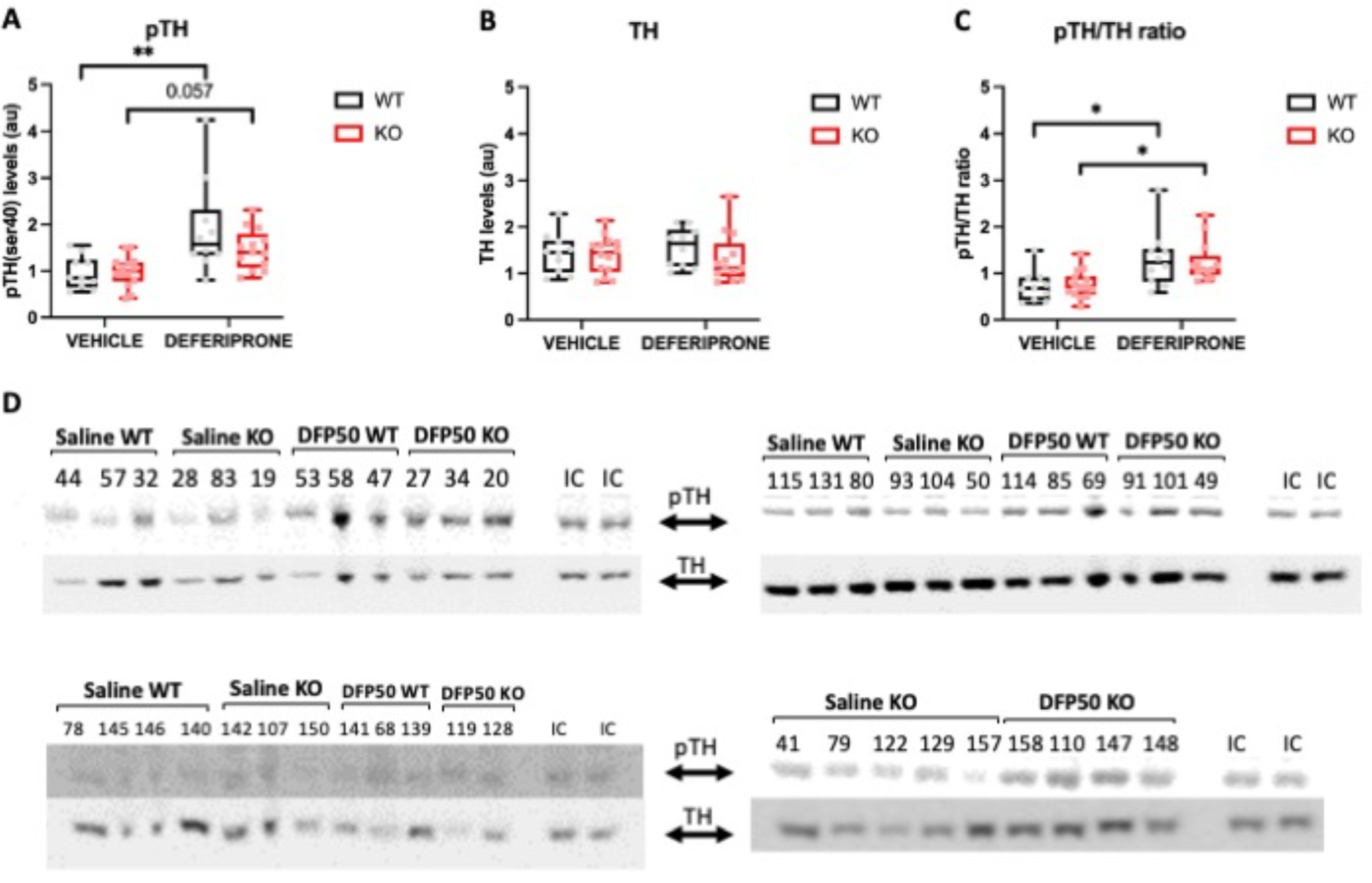
Effect of acute deferiprone treatment on pTH-Ser40, TH protein levels and the ratio of pTH-Ser40/TH in both WT and 5-HTT KO mice measured via western blot analysis. Acute deferiprone treatment increased pTH-Ser40 protein levels in WT mice as well as having a trend for increased levels in 5-HTT KO mice (A). There was no effect of acute deferiprone treatment on protein levels of TH (B). There was an increase in the ratio of pTH-Ser40/TH levels in both genotypes following acute deferiprone treatment (C). Example western blots showing the quantified band at ∼60kDa for each treatment group (D). *p<0.05; **p<0.01. n=10-14; IC = internal control

Acute deferiprone treatment resulted in an increase in levels of pTH-Ser40 in WT mice (KW=16.06, p<0.01) with a trend for an increase in 5-HTT KO mice (p=0.057) compared to vehicle treated mice in each genotype (Fig. 5A). However, there was no effect of deferiprone treatment or genotype on protein levels of TH (KW=2.64, p>0.05) (Figure 5B). Finally, there was increase in in pTH/TH ratio in both genotypes following deferiprone treatment when comparing against vehicle treated mice in both WT (KW=16.67, p<0.1) and 5-HTT KO (p<0.01) genotypes (Fig. 5C).

## 4. Discussion

Our study is the first to explore molecular actions of deferiprone following the discovery that the drug has acute antidepressant-like effects. This was done by exploring the effects of deferiprone on the phosphoproteome of the 5-HTT KO mouse model of depression. Analysing prefrontal cortex samples in WT mice, we found that deferiprone predominantly implicated phosphoproteins which are involved in cytoskeletal organisation. Meanwhile in KO mice, the phosphoproteome data indicated that proteins involved in glutamatergic signalling, synaptic, dopaminergic and cytoskeletal organisation were affected by deferiprone treatment. As we have previously found deferiprone has acute antidepressant-like properties in the Porsolt swim test (PST) in KO mice and in both genotypes in the novelty-suppressed feeding test (NSFT) (Uzungil et al., 2022), it is possible that these molecular factors may be underpinning deferiprone’s potential therapeutic properties.

The drug’s molecular actions were determined through an examination of the effect of deferiprone on phosphorylation, as it is an indirect marker of protein activity and one of the most common post-translational modifications dictating biological function (Ramazi & Zahiri, 2021). Its relevance for depression research is due to the fact that dysregulation of the phosphoproteome is observed in MDD patients, as well as other CNS disorders (Martins-de-Souza et al., 2012; Morshed et al., 2021). Furthermore, phosphoproteomics analysis has been utilised to determine the acute molecular mechanism of action of antidepressant drugs such as ketamine (Xiao et al., 2020). Our study focused on determining the effect of deferiprone in the prefrontal cortex as we had previously found that the drug increased neural activity in this region in both genotypes (Uzungil et al., 2022), in addition to the prefrontal cortex being relatively homogenous in regards to cytoarchitecture and function compared to other brain regions.

### 4.1 Acute deferiprone treatment changes phosphorylation of proteins involved in synaptic, glutamatergic and dopaminergic signalling in 5-HTT KO mice

In the 5-HTT KO deferiprone treated group, the GO analysis heavily implicated synaptic signalling, which was not affected in WT mice treated with deferiprone. Synaptic signalling is dysfunctional in MDD patients and the therapeutic properties of existing antidepressants has been linked to activity at the synapse (Bath et al., 2012; R. S. Duman et al., 2016; Kang et al., 2012; N. Li et al., 2010). 5-HTT KO mice also have deficits in synaptic plasticity and dendritic spine densities, indicating a potential critical dysfunction resulting in the observed depression-related phenotype of this mouse model (Chaji et al., 2021; Wilson et al., 2021). It is therefore possible that the 5-HTT KO selective effects of deferiprone on phosphorylation of synaptic signalling proteins may be due to genotype specific synaptic signalling dysfunction. The 5-HTT KO selective effects on protein phosphorylation may also explain the antidepressant-like behavioural effects we have previously observed in this depression mouse model (Uzungil et al., 2022). A phosphoproteomics analysis of ketamine treatment had also found effects on synaptic transmission, indicating a potential shared mechanism of action with deferiprone (Xiao et al., 2020). We have identified Pacsin1a, a phosphoprotein involved in synaptic function, as hypophosphorylated by deferiprone. Interestingly, the differentially expressed Serine 343 phosphosite of Pacsin1 has been shown to result in internalisation of AMPA receptors when phosphorylated (Anggono et al., 2013), suggesting that deferiprone may increase AMPA receptor availability by preventing internalisation.

Another synaptic protein which was hypophosphorylated following deferiprone treatment in 5-HTT KO mice was the scaffolding protein disks large homolog 4 (Dlg4), otherwise known as PSD-95, on the serine 418 phosphosite. Interestingly, this phosphosite is regulated upstream by LTP suggesting a direct link between deferiprone treatment and synaptic function in the 5-HTT KO mice (J. Li et al., 2016). Serotonin plays a modulatory role on synaptic plasticity through an interaction with synaptic density proteins including PSD-95, which may underpin genotype specific effects of deferiprone in our model of serotonergic dysfunction (Bécamel et al., 2004; Lesch & Waider, 2012).

PSD-95 also plays a key role in glutamatergic signalling, as it was 1 of 3 differentially expressed phosphosites following deferiprone treatment in 5-HTT KO mice linked to the KEGG “glutamatergic synapse” pathway. This was not surprising as synaptic processes are altered by glutamatergic neurotransmitter signalling (Popoli et al., 2011). The other 2 phosphosites implicated in “glutamatergic synapse” pathway were the transporters, Slc1a2 otherwise known as GLT-1 (which had the largest increase in fold-change) and Slc38a1. Although the GLT-1 differentially expressed phosphosite has not been previously annotated, GLT-1 has been implicated in antidepressant activity and is a glial glutamate transporter (Pham et al., 2020). Ketamine and fluoxetine’s antidepressant properties have both been linked to their actions at GLT-1 (J.-X. Chen et al., 2014; Gasull-Camós et al., 2017, 2018; Jia et al., 2020). Ketamine blocks GLT-1 activity which results in an increase in AMPA receptor activity, which therein has its antidepressant-like actions in the prefrontal cortex through an enhancement of serotonergic activity, identifying the link between GLT-1 function and serotonergic transmission (Gasull-Camós et al., 2017, 2018). The serine 52 hyper-phosphorylation of Slc38a1 has been linked to upstream regulation by RICTOR, which enhances mTOR signalling (Robitaille et al., 2013). mTOR synaptic signalling has been linked to rapid antidepressant effects of NMDA receptor antagonists such as ketamine, and may therefore signify a downstream effect on phosphorylation at Slc38a1 (N. Li et al., 2010). Therefore, the 5-HTT KO selective effect of deferiprone on glutamatergic transmission and synaptic signalling may potentially underpin the previously discovered genotype specific antidepressant-like effect in the PST (Uzungil et al., 2022).

Another protein of interest which was altered following deferiprone treatment in 5-HTT KO mice was tyrosine hydroxylase (TH), which is a rate-limiting enzyme involved in the synthesis of catecholamine neurotransmitters. This is relevant as iron is a co-factor in TH and necessitates iron availability for its function (Ramsey et al., 1996). Interestingly, the increase in phosphorylation of the serine 40 residue following acute deferiprone treatment in 5-HTT KO mice, observed in the phosphoproteomics analysis, has been implicated as a downstream effect of various antidepressant and antipsychotic drugs including ketamine, haloperidol and clozapine (Håkansson et al., 2004; Nishi et al., 2008; Snyder et al., 2007). This evidence suggests that deferiprone has a similar effect of phosphorylating TH as other psychiatric drugs, which is known to be an important mechanism for their therapeutic actions.

Our western blot analysis confirmed the increase in phosphorylation of TH of the serine 40 residue observed in 5-HTT KO mice during the phosphoproteomics analysis. Additionally, the western blot analysis also found an increase pTH-Ser40 following acute deferiprone treatment in WT mice. Importantly we observed no change in levels of TH protein due to genotype or deferiprone, indicating that the effects of deferiprone are post-translational and underpin rapid changes in protein activity rather than abundance. These findings confirm previous observations in cell culture that iron chelators such as deferoxamine increase levels of pTH and the pTH/TH ratio, which was reflected in our deferiprone findings (Connor et al., 2009). Phosphorylation of the serine 40 residue of TH has been shown to play a critical role in dopaminergic transmission, synaptic plasticity and reward related behavioural outcomes, factors that are all implicated in MDD (Liu et al., 2018; McIntosh et al., 2013). In addition to glutamatergic and synaptic protein phosphorylation effects of acute deferiprone treatment which we have mentioned, TH and dopaminergic pathways might be an additional molecular mechanism which contributes to the antidepressant-like properties of deferiprone. Critically, linking these mechanisms, dopaminergic pathways are important for ketamine’s antidepressant properties, suggesting a combinatory role of glutamatergic and dopaminergic transmission in effective antidepressants (Belujon et al., 2016; Belujon & Grace, 2017; Carreno et al., 2016). These findings suggest that acute iron chelation increases phosphorylation of TH at the serine 40 reside which via downstream pathways has the potential to have positive effects on behaviours associated with MDD.

It is possible that the acute antidepressant/anxiolytic-like effects of deferiprone observed in the NSFT in both genotypes is primarily driven by an increase in pTH-Ser40 levels (Uzungil et al., 2022). This hypothesis would be in line with prior findings that reward processes and dopaminergic signalling are implicated in NSFT performance (Bahi & Dreyer, 2019; Shuto et al., 2020). Moreover, the lateral amygdala is also involved in consummatory motivational processing in the NSFT (Cabeza et al., 2021) and was a region which we found to be heavily implicated in the action of deferiprone (Uzungil et al., 2022), Therefore, it is possible that the behavioural effects of deferiprone in both genotypes in the NSFT is driven by the increase in pTH-Ser40 levels.

Although our western blot analysis reflects the findings of increased pTH-Ser40 expression in 5-HTT KO mice during phosphoproteomics analysis, the increase in WT mice in western blot analysis was not replicated in the phosphoproteomics analysis. It is possible that the increased variability in WT groups compared to 5-HTT KO groups due to batch effects as observed on the PCA plot is resulting in an inability to detect all differentially expressed phosphopeptides and resulting in false negatives in our analysis. Future experimentally controlled phosphoproteomics experiments should be conducted to confirm these findings.

### 4.2 Acute deferiprone treatment effects phosphorylation of proteins involved in cytoskeletal organisation in both genotypes

The phosphoproteomics dataset indicates that in both genotypes, cytoskeletal organisation signalling was affected by acute deferiprone treatment in the prefrontal cortex. A prominent cytoskeletal protein which was the only hypophosphorylated phosphopeptide following deferiprone treatment in both genotypes was alpha-adducin (ADD1). Another adducin protein beta-adducin (ADD2), which had the second largest fold-change increase following deferiprone treatment in 5-HTT KO mice, has been shown to be altered in the brains of patients with MDD (Martins-de-Souza et al., 2012). Furthermore, hyperphosphorylation of the ADD2 Serine 602 phosphosite has been linked to upstream regulation by ERK 1/2 activity (Zennadi et al., 2012), which is a signalling pathway involved in the antidepressant-like effects of ketamine (Réus et al., 2014). Neuronal plasticity is dependent upon the dynamics of the cellular cytoskeleton to alter the architecture of the dendrite and form neurite outgrowths (Fukazawa et al., 2003; Szabó et al., 2016). Post-translational modification of cytoskeletal proteins have been shown to be maladaptive in MDD and stress induced models of depression (Martins-de-Souza et al., 2012; Soetanto et al., 2010; Wong et al., 2013). Conversely, both fluoxetine and ketamine have actions on cytoskeletal proteins to alter synaptic remodelling, necessary for their antidepressant-like actions (Bianchi et al., 2009; Colic et al., 2019; Zunszain et al., 2013). One example of a protein which was differentially expressed following deferiprone treatment in WT mice was stathmin 1 (STMN1) which has a role in microtubule dynamics and plays a role in synaptic plasticity. Polymorphisms of the *STMN1* gene in humans has been linked to cognitive and affective disturbances indicating the role of this protein in depression-related behaviours including emotional interference processing (Ehlis et al., 2011). The KEGG pathway analysis revealed that STMN1 also plays a role in the MAPK signalling pathway, which is implicated in the rapid antidepressant-like effects of ketamine in animal models (Réus et al., 2014). The shared effects of deferiprone on cytoskeletal proteins in both WT and 5-HTT KO mice may explain the antidepressant-/anxiolytic-like effect of the drug in the NSFT in both genotypes (Uzungil et al., 2022). Finally, in agreement with our previous findings that deferiprone was not altering iron levels in various brain regions following acute treatment (Uzungil et al., 2022), we found that there were only a couple of phosphopeptides involved in iron regulation (BLVRA and NDRG1). Our findings indicate that deferiprone is predominantly acting in an iron-independent manner to phosphorylate proteins in the prefrontal cortex. This would suggest that deferiprone is acting in an iron-independent manner to exert its antidepressant-like properties.

### 4.3 Acute deferiprone treatment phosphorylates proteins involved in NOS signalling

PSD-95 function is altered in the prefrontal cortex in MDD and it is theorised to be a novel avenue for antidepressant treatment, through its interactions with NMDA receptors and nNOS complex (Doucet et al., 2012; Feyissa et al., 2009; Karolewicz et al., 2009). As deferiprone has been shown to effect eNOS activity peripherally, the PSD-95/NMDA/nNOS complex may be implicated in the actions of the drug (Doucet et al., 2012; Sriwantana et al., 2018). Further evidence for the potential role of the NOS pathway in the actions of deferiprone is reflected in the hypophosphorylation of a phosphosite of the histamine H1 receptor (HRH1) in 5-HTT KO mice, as histamine has been shown to increase nNOS activity at HRH1, however the serine 344 phosphosite has not been extensively annotated (Yang & Hatton, 2002). In addition to HRH1, in WT mice we found that deferiprone treatment resulted in reduced phoshphorylation of serine 2 phosphosite on the NDRG1 protein. Nitrosative stress, which is implicated in MDD, antidepressants and iron regulation, is involved in NDRG1 activity whereby nitric oxide upregulates NDRG1 activity (Dhir & Kulkarni, 2011; Finkel et al., 1996; Hickok et al., 2011; Lee et al., 2006; Suzuki et al., 2001). Iron chelation by non-deferiprone drugs including deferoxamine were shown to upregulate NDRG1 levels, which influences neurotransmitter release, however the effect on this phosphosite was not explored (Kovacevic & Richardson, 2006; Le & Richardson, 2004). The effect of deferiprone on NOS signalling proteins PSD-95, HRH1 and NDRG1 in either genotype coupled with role of iron chelators in NOS pathways demands further exploration of the potential NOS-driven acute antidepressant-like properties of deferiprone.

### 4.4 No substantial effect to phosphoproteome or global proteome due to genetic ablation of 5-HTT

We also examined the effect of 5-HTT genetic ablation on the phosphoproteome in the prefrontal cortex in the absence of treatment and found a smaller number of differentially expressed phosphosites. The GO overrepresentation analysis did not reveal any process as being increased in the differential expression list against a reference genome indicating there was not a large change to the phosphoproteome. However, many kinases and phosphatases have been previously implicated in 5-HTT trafficking and function which we did not find to have an effect on the phosphoproteome in the prefrontal cortex (Annamalai et al., 2012; Goodfellow et al., 2014; Jayanthi et al., 2005; Ragu Varman et al., 2021; Samuvel et al., 2005). Although the prefrontal cortex is dysregulated in 5-HTT KO mice, it may not be the region of the most widespread changes in kinase/phosphatase activity. Instead, regions which we found to be differentially activated in 5-HTT KO mice in our previous c-Fos study included the dorsal raphe and amygdala (Uzungil et al., 2022), which may exhibit more extensive changes to the phosphoproteome. Furthermore, the hippocampus is another region which has extensive changes in the 5-HTT KO mice and may be another region of interest (Renoir et al., 2008; Wilson et al., 2021).

We were also able to confirm that the detected differential expression of phosphosites were not due to changes in total protein levels as none of the targets were altered in expression in the global proteomics analysis. Proteins such as BNDF and synaptic scaffolding proteins from previous studies that were shown to be reduced in KO rodents unfortunately were not identified in the current study (Brivio et al., 2019; Molteni et al., 2010). This is likely due to the methodology of proteomics analysis which is not sensitive to proteins with low abundance (Geiger et al., 2012; Nguyen et al., 2019). However, as proteins associated with our differentially expressed phosphosites were not changed in the global proteomics analysis, this indicates that the phosphoproteomics results were not confounded by changes in absolute protein levels in 5-HTT KO mice.

An interesting finding was that compared to 5-HTT KO mice, WT mice treated with deferiprone had greater proportion of differentially expressed phosphosites which were hypophosphorylated. Compared to 5-HTT KO mice this would suggest that deferiprone is increasing phosphatase activity or reducing the activity of kinases. As none of the differentially expressed phosphosites following deferiprone treatment were differentially expressed constitutively due to genetic ablation of the 5-HTT either during phosphoproteomics analysis, or the associated protein in global proteomics analysis, these findings would suggest a differential molecular response to deferiprone treatment in each genotype. This finding would be compatible with our previous reporting on the difference in behavioural, brain region activity and functional network connectivity following deferiprone treatment in either genotype (Uzungil et al., 2022). It is therefore possible that the 5-HTT KO specific therapeutic properties of deferiprone may be driven by an increase in phosphorylation/activity of proteins implicated in depression pathways which is not replicated in WT animals.

### 4.5 Limitations and conclusions

Complex brain regions such as the prefrontal cortex are likely to contain heterogeneous cell types which have the potential to mask phosphorylation effects of the drug relevant to the therapeutic mechanism. A phosphoproteomics analysis of smaller regions with cellular homogeneity may reveal more robust effects of deferiprone on molecular targets. As deferiprone has been previously shown to have antidepressant-like properties and the current phosphoproteomics analysis has identified molecular mechanisms involved in the action of existing antidepressant, this study is an important benchmark for further experimentation. It would also be informative to examine other brain regions implicated in the antidepressant-like properties of deferiprone and MDD such as the amygdala, as they may reveal additional molecular factors involved in deferiprone’s therapeutic mechanism of action.

The data indicates that acute deferiprone treatment on the prefrontal cortex alters phosphorylation of proteins involved in synaptic, glutamatergic and dopaminergic signalling in the 5-HTT KO mouse model of depression. Additionally, there was an effect on proteins involved in cytoskeletal organisation irrespective of genotype. It is likely too simplistic to suggest that one protein or neurotransmitter is involved in the positive effects of a therapeutic in a complex disorder like MDD. Instead, it is probable that a combinatory effect of glutamate and dopamine neurotransmitter signalling on synaptic plasticity and cytoskeletal protein mechanisms are involved in deferiprone’s antidepressant-like properties. The genotype specific effect of deferiprone on the 5-HTT KO mouse may be driven by the intrinsic synaptic and glutamatergic signalling pathology and could be responsible for the genotype specific antidepressant-like properties of the drug. These results would indicate that deferiprone has actions similar to the fast-acting antidepressant glutamatergic antagonist ketamine. While our study reveals multiple promising targets, future analysis of these phosphopeptides is necessary to pinpoint the critical proteins and signalling pathways involved in deferiprone’s therapeutic actions, especially in regions heavily implicated in MDD.

## Acknowledgements

This work was supported by a National Health and Medical Research Council (NHMRC) Project Grant. TR is a NHMRC Boosting Dementia Research Leadership Fellow. AJH is a NHMRC Principal Research Fellow. The Florey Institute of Neuroscience and Mental Health (FINMH) acknowledges the support from the Victorian Government’s Operational Infrastructure Support Grant.

