## Supplementary figures and images for "Phosphoproteomics implicates glutamatergic and dopaminergic signalling in the antidepressant-like properties of the iron chelator deferiprone"

### Supplementary Figure 1

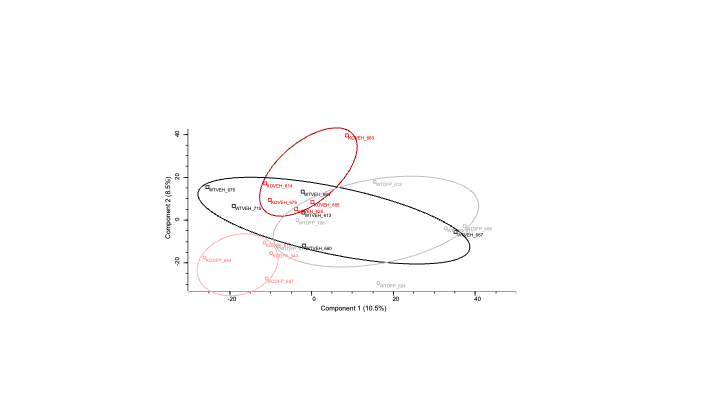

### Supplementary Figure 2

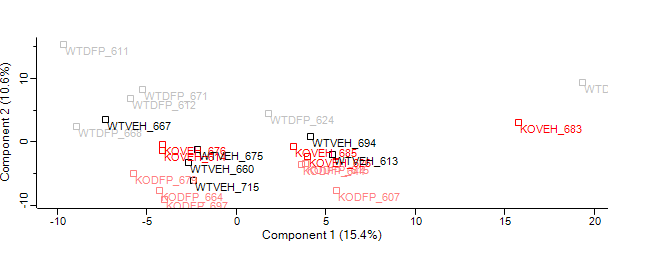

### Supplementary Figure 3

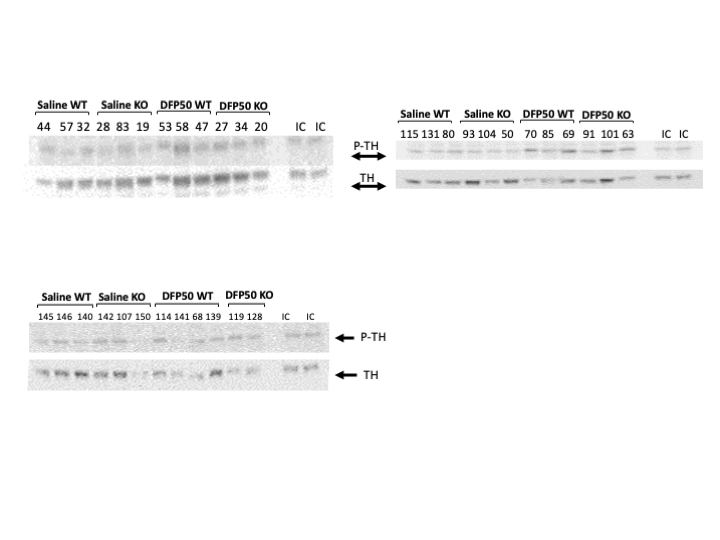
